# Multiple bouts of high-intensity interval exercise reverse age-related functional connectivity disruptions without affecting motor learning in older adults

**DOI:** 10.1101/2021.03.07.434269

**Authors:** Brian Greeley, Briana Chau, Christina B. Jones, Jason L. Neva, Sarah N. Kraeutner, Kristin L. Campbell, Lara A. Boyd

**Affiliations:** Department of Physical Therapy, University of British Columbia, Vancouver, Canada, V6T1Z3; Graduate Program in Rehabilitation Sciences, University of British Columbia, Vancouver, Canada, V6T1Z3; UBC MRI Research Centre, University of British Columbia, Vancouver, Canada, V6T2B5; Université de Montréal, École de kinésiologie et des sciences de l’activité physique, Faculté de médecine, Montréal, QC, Canada, H3C3T5; Centre de recherche de l’institut universitaire de gériatrie de Montréal, Montréal, QC, Canada, H3T1P1

## Abstract

Exercise has emerged as an intervention that may mitigate age-related resting state functional connectivity and sensorimotor declines. Here, 42 healthy older adults rested or completed 3 sets of high-intensity interval exercise for a total of 23 minutes, then immediately practiced an implicit motor task with their non-dominant hand across five separate sessions. Participants completed resting stage functional MRI before the first and after the fifth day of practice; they also returned 24-hours and 35-days later to complete short- and long-term retention. Independent component analysis of resting state functional MRI revealed increased connectivity in the frontoparietal, the dorsal attentional, and cerebellar networks and group. seed-based analysis showed strengthened connectivity between the limbic system and right cerebellum, and between the right cerebellum and bilateral middle temporal gyri in the exercise group. There was no motor learning advantage for the exercise group. Our data suggest that exercise paired with an implicit motor learning task in older adults can augment resting state functional connectivity without enhancing behaviour beyond that stimulated by skilled motor practice.

**Impact statement:** Five separate bouts of exercise paired with skilled motor practice strengthens resting state networks in brain regions that are susceptible to declines in older adults without affecting motor acquisition or learning.

## Introduction

The global population is aging^1^. Given that sensorimotor decline accompanies aging ^2,3^, there is growing interest in interventions that mitigate these changes in older adults. Data from studies of young adults suggest that exercise can enhance cortical excitability and may improve motor learning. Following a single bout of exercise, transcranial magnetic stimulation measures show changes in intracortical circuit excitability ^4,5^ in young adults. Similarly, 20 minutes of high-intensity interval exercise, determined by a high percentage (75-90%) of workload taken from a separate maximal exercise stress test, preceding practice enhances motor learning in young adults, regardless of the motor task. ^6–8^. However, the majority of research investigating the impact of high-intensity interval exercise has focused on healthy young individuals; no previous work has considered whether pairing skilled motor practice with high-intensity interval exercise will impact motor learning in older adults.

As the limbic system is impacted by both chronic ^9,10^ and single, 20 min bouts of exercise ^11^, motor learning tasks that engage these brain regions may also be enhanced following exercise. For example, 3 days of aerobic exercise per week over the course of 1 year, increases the size of the hippocampus, which leads to improvements in spatial memory in older adults ^9^. A month of exercise enhances learning along with hippocampal neurogenesis in aged mice ^12^, and is neuroprotective in both the hippocampus and amygdala in young mice ^13,14^. Functional magnetic resonance imaging (fMRI) studies also support the role of the limbic system in learning, with the hippocampi engaged in both explicit and implicit motor learning in older adults ^15^. Taken together, the limbic system appears to be positively impacted by aerobic exercise which in turn may facilitate motor learning.

Exercise may mitigate age-related decreases in resting state functional connectivity (FC), but whether changes in FC are accompanied by motor learning in older adults is unknown. Resting state FC is a neuroimaging method that captures the temporal coherence of spatially distinct brain regions at rest. Age-related decreases in FC networks have been well documented, especially in the default mode network (DMN)^16–20^. Moreover, age-related decreases in FC may have functional significance; the weaker the FC within the DMN relates to poorer executive function, slower processing speed ^17^ and poorer memory performance ^20,21^. It is possible that exercise mitigates age-related decreases within the DMN. For example, Weng et al. (2016) found that older adults displayed increases in FC between the hippocampus and the medial prefrontal cortex immediately following a 30 minute bout of exercise, and Voss et al. (2010) reported an increase in FC between the posterior cingulate cortex and frontal medial cortex and the posterior cingulate cortex and middle temporal gyrus after 6 months of a walking intervention. Importantly, Voss et al. (2010) found no benefit of exercise on a battery of cognitive measures in older adults, suggesting that exercise may have the capacity to mitigate age-related FC disruptions, especially for brain regions within the DMN but may not always affect behaviour.

In the current study, healthy older adults practiced an implicit motor sequence task following a bout of high-intensity interval exercise in five separate sessions. Additionally, participants completed MRI scans before and after the intervention. We hypothesized that the participants in the exercise group would show greater change in both motor skill acquisition and motor learning at the retention tests compared to the rest group. We also hypothesized that those in the exercise, but not the rest group, would display increases in connectivity in networks and brain regions typically disrupted with age including the DMN and limbic brain regions areas.

## Methods

### Participants

Forty-four participants were recruited from Vancouver, British Columbia or the greater Vancouver area. Twenty were randomly assigned and to the exercise group (9 males; mean age 68.3 years old), whereas 22 participants were randomized into the rest group (5 males; mean age 65.2 years old). We recruited both left and right handed individuals; handedness was determined using the Edinburgh Handedness Scale ^22^. Demographic information was collected verbally during the first session. Usual physical activity levels were assessed using the Godin Leisure Time Physical Activity questionnaire ^23^ and collected during the first session (Table 1). Participants were healthy with no reported neurological disorders and screened for any potential contraindications to magnetic resonance imaging using standard screening forms. The Clinical Research Ethics Board at the University of British Columbia approved all protocols, and all methods were performed in accordance with relevant guidelines and regulations.

**Table 1.**
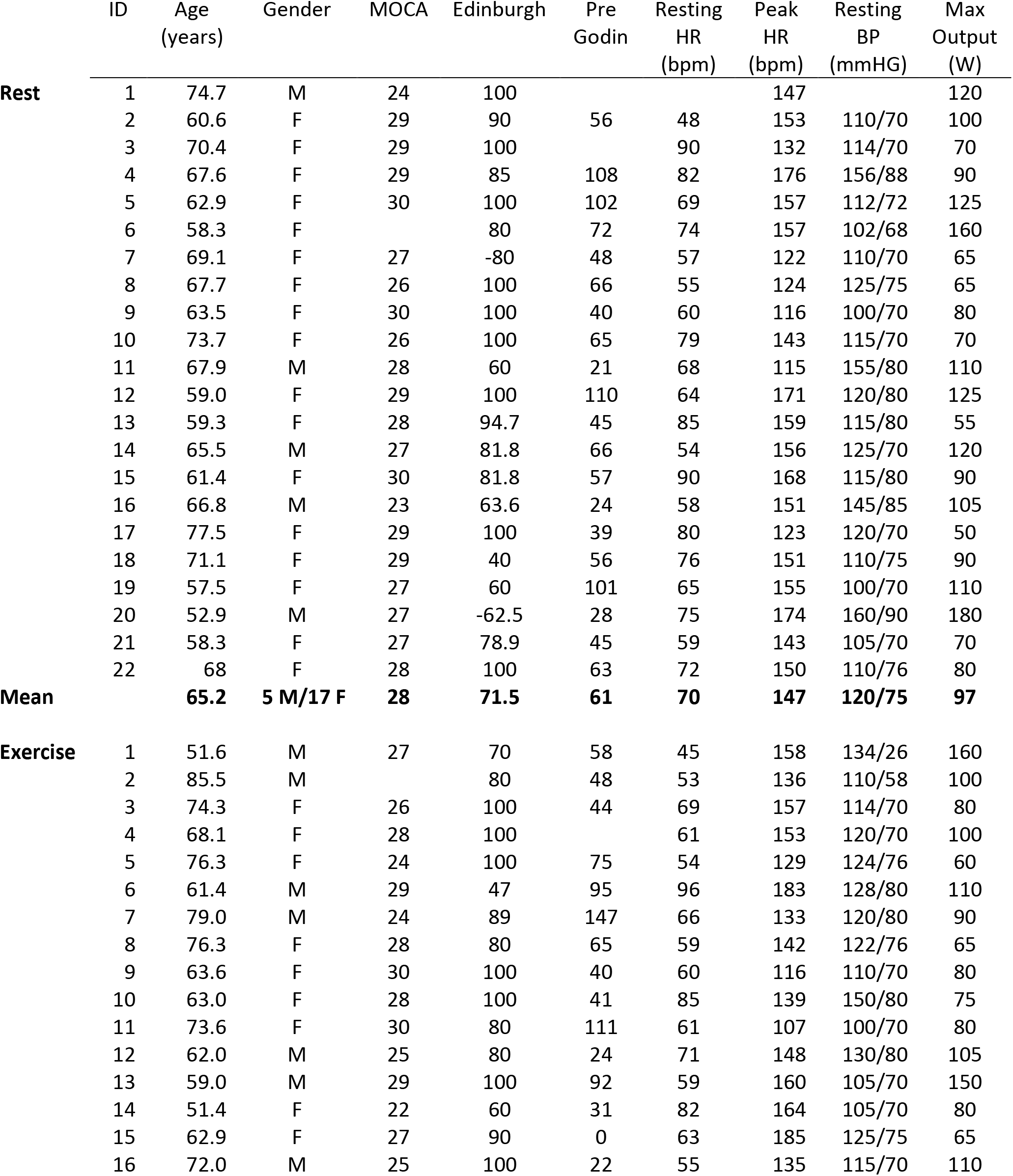
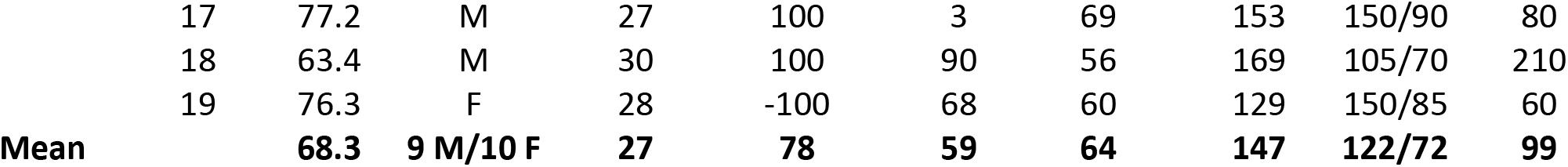
Participant demographic table. Resting heart rate (HR), peak HR, resting blood pressure (BP), and max output in watts (W) are values taken from the stress test.

|  | ID | Age<br>(years) | Gender | MOCA | Edinburgh | Pre<br>Godin | Resting<br>HR<br>(bpm) | Peak<br>HR<br>(bpm) | Resting<br>BP<br>(mmHG) | Max<br>Output<br>(W) |
| --- | --- | --- | --- | --- | --- | --- | --- | --- | --- | --- |
| **Rest** | 1 | 74.7 | M | 24 | 100 |  |  | 147 |  | 120 |
|  | 2 | 60.6 | F | 29 | 90 | 56 | 48 | 153 | 110/70 | 100 |
|  | 3 | 70.4 | F | 29 | 100 |  | 90 | 132 | 114/70 | 70 |
|  | 4 | 67.6 | F | 29 | 85 | 108 | 82 | 176 | 156/88 | 90 |
|  | 5 | 62.9 | F | 30 | 100 | 102 | 69 | 157 | 112/72 | 125 |
|  | 6 | 58.3 | F |  | 80 | 72 | 74 | 157 | 102/68 | 160 |
|  | 7 | 69.1 | F | 27 | -80 | 48 | 57 | 122 | 110/70 | 65 |
|  | 8 | 67.7 | F | 26 | 100 | 66 | 55 | 124 | 125/75 | 65 |
|  | 9 | 63.5 | F | 30 | 100 | 40 | 60 | 116 | 100/70 | 80 |
|  | 10 | 73.7 | F | 26 | 100 | 65 | 79 | 143 | 115/70 | 70 |
|  | 11 | 67.9 | M | 28 | 60 | 21 | 68 | 115 | 155/80 | 110 |
|  | 12 | 59.0 | F | 29 | 100 | 110 | 64 | 171 | 120/80 | 125 |
|  | 13 | 59.3 | F | 28 | 94.7 | 45 | 85 | 159 | 115/80 | 55 |
|  | 14 | 65.5 | M | 27 | 81.8 | 66 | 54 | 156 | 125/70 | 120 |
|  | 15 | 61.4 | F | 30 | 81.8 | 57 | 90 | 168 | 115/80 | 90 |
|  | 16 | 66.8 | M | 23 | 63.6 | 24 | 58 | 151 | 145/85 | 105 |
|  | 17 | 77.5 | F | 29 | 100 | 39 | 80 | 123 | 120/70 | 50 |
|  | 18 | 71.1 | F | 29 | 40 | 56 | 76 | 151 | 110/75 | 90 |
|  | 19 | 57.5 | F | 27 | 60 | 101 | 65 | 155 | 100/70 | 110 |
|  | 20 | 52.9 | M | 27 | -62.5 | 28 | 75 | 174 | 160/90 | 180 |
|  | 21 | 58.3 | F | 27 | 78.9 | 45 | 59 | 143 | 105/70 | 70 |
|  | 22 | 68 | F | 28 | 100 | 63 | 72 | 150 | 110/76 | 80 |
| **Mean** |  | **65.2** | **5 M/17 F** | **28** | **71.5** | **61** | **70** | **147** | **120/75** | **97** |
| **Exercise** | 1 | 51.6 | M | 27 | 70 | 58 | 45 | 158 | 134/26 | 160 |
|  | 2 | 85.5 | M |  | 80 | 48 | 53 | 136 | 110/58 | 100 |
|  | 3 | 74.3 | F | 26 | 100 | 44 | 69 | 157 | 114/70 | 80 |
|  | 4 | 68.1 | F | 28 | 100 |  | 61 | 153 | 120/70 | 100 |
|  | 5 | 76.3 | F | 24 | 100 | 75 | 54 | 129 | 124/76 | 60 |
|  | 6 | 61.4 | M | 29 | 47 | 95 | 96 | 183 | 128/80 | 110 |
|  | 7 | 79.0 | M | 24 | 89 | 147 | 66 | 133 | 120/80 | 90 |
|  | 8 | 76.3 | F | 28 | 80 | 65 | 59 | 142 | 122/76 | 65 |
|  | 9 | 63.6 | F | 30 | 100 | 40 | 60 | 116 | 110/70 | 80 |
|  | 10 | 63.0 | F | 28 | 100 | 41 | 85 | 139 | 150/80 | 75 |
|  | 11 | 73.6 | F | 30 | 80 | 111 | 61 | 107 | 100/70 | 80 |
|  | 12 | 62.0 | M | 25 | 80 | 24 | 71 | 148 | 130/80 | 105 |
|  | 13 | 59.0 | M | 29 | 100 | 92 | 59 | 160 | 105/70 | 150 |
|  | 14 | 51.4 | F | 22 | 60 | 31 | 82 | 164 | 105/70 | 80 |
|  | 15 | 62.9 | F | 27 | 90 | 0 | 63 | 185 | 125/75 | 65 |
|  | 16 | 72.0 | M | 25 | 100 | 22 | 55 | 135 | 115/70 | 110 |
| Running head: EXERCISE AFFECTS CONNECTIVITY IN ELDERLY |  |  |  |  |  |  |  |  |  |  |
|  | 17 | 77.2 | M | 27 | 100 | 3 | 69 | 153 | 150/90 | 80 |
|  | 18 | 63.4 | M | 30 | 100 | 90 | 56 | 169 | 105/70 | 210 |
|  | 19 | 76.3 | F | 28 | -100 | 68 | 60 | 129 | 150/85 | 60 |
| **Mean** |  | **68.3** | **9 M/10 F** | **27** | **78** | **59** | **64** | **147** | **122/72** | **99** |

### Study Design

Participants reported to the laboratory for a total of 11 sessions. During the first session, participants provided consent and completed a cardiologist supervised maximally graded stress test. All participants, regardless of group assignment, were cleared for exercise by a licensed cardiologist. In session 2, participants underwent a baseline MRI that included T1 anatomy and resting state scans. Participants were then pseudo-randomized, accounting for age and sex using a custom computer program, to “exercise” or “rest” conditions. In sessions 3 and 10 neurophysiology was assessed using transcranial magnetic stimulation (unpublished). In sessions 4 through 8 participants either watched a nature documentary or exercised for 23 minutes before completing 4 blocks of a complex, implicit motor sequence learning task (see serial targeting task below). To allow enough time in between each exercise session to enable muscle recovery, we scheduled approximately two practice sessions per week, which is a feasible recommendation and in line with current aerobic exercise guidelines ^24^. Session 9 contained a 24-hour retention session where participants completed 1 block of the motor task to assess short-term motor learning and underwent their second resting state scan (post-intervention scan). Thirty-five days after session 9, participants returned to complete a retention test composed of 1 block of the motor task to assess long-term motor learning (Figure 1). The purpose of the 35-day retention session was to assess the long-term impact and time course of exercise on motor learning.

**Figure 1.**
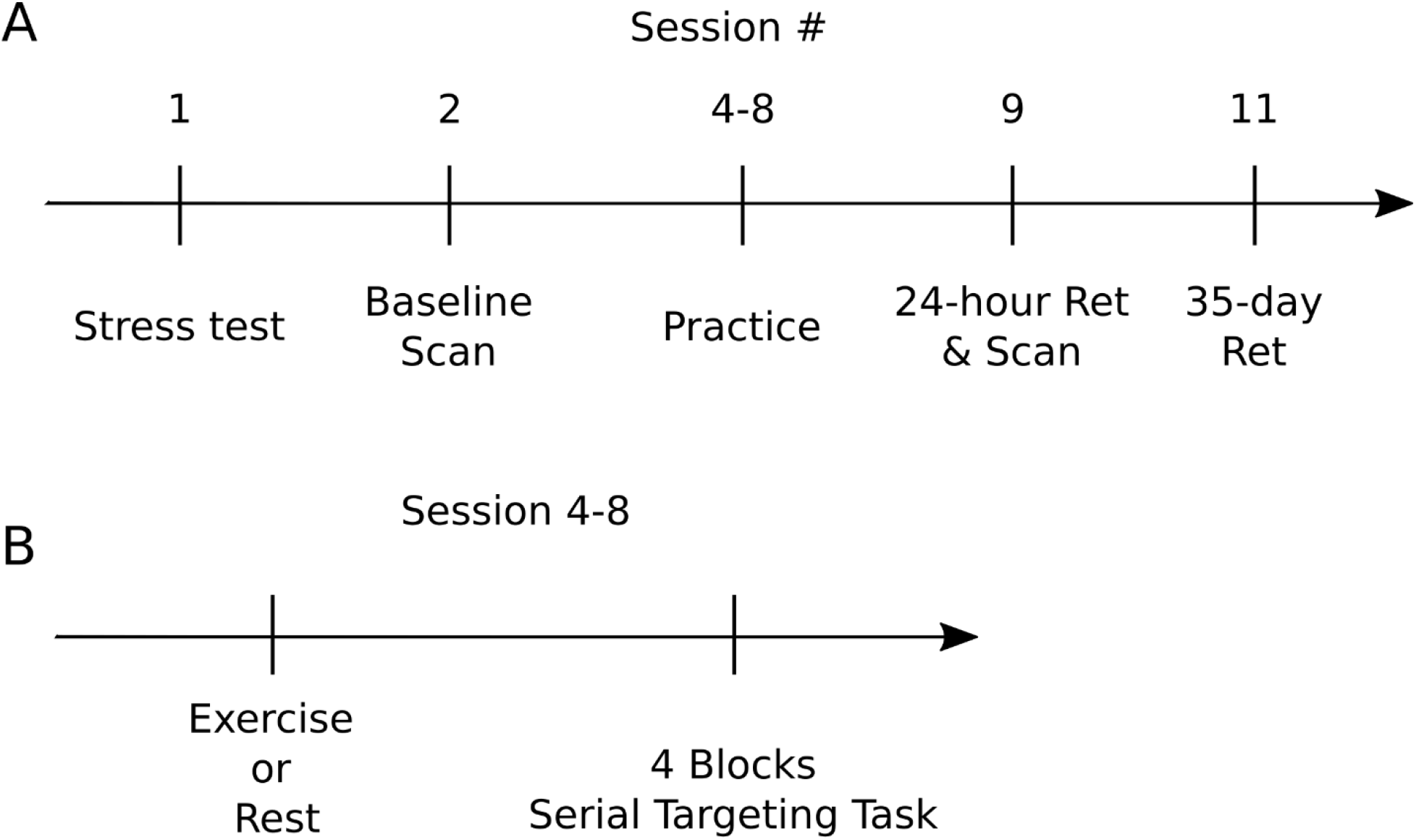
Experimental design of: (A) the entire study and (B) an exercise or rest practice session. Regardless of group, all participants completed a maximally graded stress in session 1. Session 2 and 9 contained pre- and post-intervention MRI resting state scans, respectively. In sessions 4-8 participants either watched 23 minutes of a nature documentary or underwent 23 minutes of high intensity exercise before practicing the implicit motor task. In addition to the post-intervention MRI scan, session 9 contained the 24-hour, short-term retention test to assess motor learning. Session 11 included the 35 day, long-term retention test.

### Stress Test

All participants completed a cardiologist supervised maximally graded stress test during their first visit. In addition to ensuring that all participants were safe to exercise, data from the stress test were used to determine the participant’s high-intensity workload for training sessions. Before attending the stress test, participants were instructed to refrain from engaging in vigorous physical activity for 24 hours prior and not to eat a large meal 2 hours before to their scheduled test. Once participants arrived, they provided informed consent, and a list of current medications and past medical history (if applicable). Electrocardiogram leads were then placed on the participant’s chest which was wirelessly transmitted to, and saved on, a computer. After approximately 3 minutes resting in supine, heart rate and blood pressure were recorded. Next, participants were seated on a recumbent bike (SciFit). The participant’s feet were secured used two Velcro straps and the seat distance and height was adjusted for each participant, so that the knee was flexed approximately between 25-45° during full leg extension and that the participant was comfortable during cycling. Heart rate was monitored continuously throughout the stress test and recorded once every minute. Blood pressure was taken manually by a cardiology technologist every other minute. Rate of perceived exertion (RPE) ^25^ was obtained once every minute. Participants were instructed to generate their power from their legs, keep their arms and hands relaxed, and maintain between 50 and 80 revolutions per minute (RPM). The stress test began with a 2-minute warm-up at 10 watts. Then resistance was increased every minute thereafter, Depending on the last reported RPE, heart rate, and blood pressure, wattage was increased either by 5, 10, or 15 based on an individual basis and current stress test recommendations ^26^. The stress test was terminated if the participant stopped pedaling or if RPMs dropped below 50 and continued to decline for more than five seconds. RPE at test termination was collected. After the termination of the stress test, participants cooled down between 2 and 3 minutes at 10 watts. After the cool down, participants laid down and rested between 1 and 5 minutes until heart rate and blood pressure returned to baseline.

### Intervention

#### Exercise

For each practice session, participants in the exercise group completed a bout of high-intensity interval exercise on a stationary recumbent bicycle (SciFit). An exercise session consisted of a warm-up and three equal repetitions of high-intensity exercise interleaved with active recovery. Participants were asked to pedal at 50 to 80 RPM throughout the session. First, participants completed a five-minute warm-up, cycling at 10 Watts. Following warm-up, participant completed 3 sets of 3-minute high-intensity bouts of exercise (at 75% of max power output as determined from the stress test), interspersed with 3-minutes of active recovery (at 10 Watts) ^6,27^. Heart rate was monitored using the Alpha 53p watch (Mio, Portland) and recorded at the last minute of every interval: namely at the end of warm up (5 minutes), first high-intensity bout (8 minutes), first active recovery bout (11 minutes), second high-intensity bout (14 minutes), second active recovery bout (17 minutes), the third high-intensity bout (20 minutes), and last active recovery bout (23 minutes).

### Rest

Participants assigned to the rest group sat for 23 minutes while watching a nature documentary (Planet Earth) during each practice session (equivalent to total duration to complete the warm and interval sessions). Heart rate was recorded at the same time intervals as the exercise group.

### Serial Targeting Task

Skilled motor practice took place using a KINARM end-point robot (BKIN Technologies Ltd, Kingston, ON, Canada). Participants were seated in the KINARM chair which was adjusted so that the individual’s head was positioned in the center of the visual field. Participants grasped one of the end-point handles using their non-dominant hand (Figure 2A). The non-dominant arm was used to increase the overall difficultly of the task. A bib was placed over the front of the participant which occluded vision of their arm and hand.

**Figure 2.**
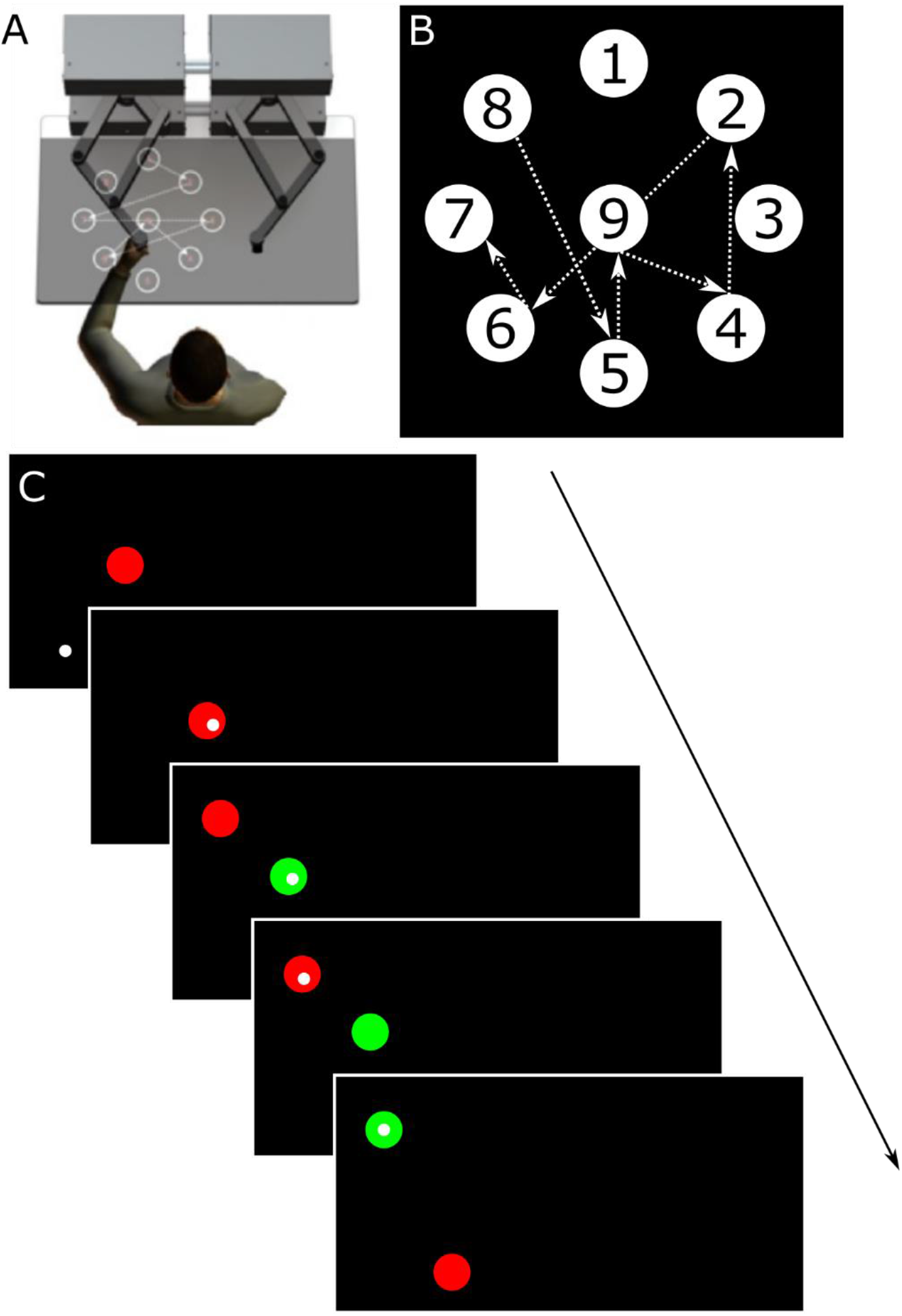
Set-up and schematic overview of the serial targeting task. The serial targeting task was practiced on A) an endpoint KINARM robot immediately following 23 minutes of rest or high intensity exercise. Participants used their non-dominant arm to control a frictionless lever to move to illuminated targets as quickly and accurately as possible. B) Numbered targets displayed represent all targets in the workspace and the order of the embeded repeated target sequence. Note, participants only saw 2 targets at any given time (when not next in the sequence the targets were not displayed) and targets were *not* numbered during the task. C) The task began with the appearance of a red center target with a small white circle representing the participant’s non-dominant hand position in real time (C, top). The red target stayed on the screen until the participant moved and held the white dot (i.e., their hand) in the red target for 500 ms, after which a new red target appeared and the current red targets turned green simultaneously. This process repeated until participants completed one block (a total of 111 movements, 48 of which belong to a 6-element repeated target sequence). A total of 4 blocks were completed during each practice session. All targets are presented on the non-dominant side of the participant’s workspace.

Motor performance and learning were assessed using the Serial Targeting Task (STT)^6,28,29^, an implicit motor sequence task. Targets were displayed on the participant’s non-dominant side of the KINARM workspace (Figure 2A). In the task there were a total of nine targets. One center target, and eight peripheral targets that formed an equidistant circular array of a 10 cm radius. Each circular target had a radius of 1.14 cm (Figure 2B).

To begin the task, participants used their non-dominant arm to reach to a red center target. To initiate the appearance of the next target, participants had to hold a white cursor, which represented the position of their hand, within the current red target for 500 ms, after which the current target turned green and simultaneously the next red target appeared (Figure 2C). Participants had 10,000 ms to reach to each target. Cursor position was sampled at 1000 Hz. Before each practice session, participants were instructed to move to each target as quickly and accurately as possible, making sure not to over- or undershoot the target. On practice day 1, participants were briefly exposed to the task (pre-test) which contained 20 trials; this took approximately 30 seconds. On each of the 5 training days, participants performed 4 blocks, or 444 movements, immediately following exercise or rest (according to group assignment). Motor learning was assessed both in the short- and long-term by two no-exercise or no-rest retention tests made up of 1 block of the STT. Short-term retention was tested 24-hours after the last day of practice and long-term retention was assessed 35 days after the last day of practice.

Unbeknownst to the participants, the STT contained a repeated sequence which was embedded between sequences of random movements. We included both random and repeated sequences to distinguish between motor learning (repeated sequences) and motor performance (random sequences) as well as keep learning implicit. Including random sequences and contrasting the two provides a specific measure of skill and reduces contamination influences of factors like stimulus-response mapping, fatigue and motivation ^30^. Also, the removal of random sequences would have exposed the nature of the task and have likely resulted in explicit learning and would have allowed participants to anticipate the targets, which may have resulted in a ceiling effect in reaction time. Participants were exposed to the same, 6-element sequence across practice sessions. Behavioural change in motor performance of repeated sequences reflects implicit motor learning whereas change in random sequence performance reflects motor control ^6,31^. Thus, participants were exposed to a repeating 6-element sequence, which was flanked by 7 targets that appeared at random. The order of the 6-element repeating sequence was: 8, 5, 9, 4, 2, 6, 7 (Figure 2B). Random sequences were never repeated but appeared in the same order across practice for all participants. Within any one block of practice, and regardless of the practice session, repeating sequence trials were always displayed in the following trial order: trial 8-13, 21-26, 34-39, 47-52, 60-65, 73-78, 86-91, 99-104. Thus, participants were exposed to 32 repetitions of the repeating sequence within one practice session (8 times per block, 4 blocks of practice; 192 trials). Within any block of practice, random sequence trials were always displayed in the following trials: 1-7, 14-20, 27-33, 40-46, 53-59, 66-72, 79-85, 92-98, 105-111. Thus, participants were exposed to 36 random sequences within a single practice session.

### Explicit Awareness

Explicit awareness was assessed during short- and long-motor learning sessions. Following the 24-hour retention test, participants were shown a series of 10 sequence movies displayed on the KINARM screen and asked to decide if they recognized any as the repeated pattern that they practiced. Three of the 10 were “true” repeating sequences (same as the repeated practice pattern); 7 were foils. Individuals who identified both the repeated and random sequences at a better than chance (i.e., 2 of 3 repeated sequences identified correctly and 4 of 7 novel, random sequences identified as not having been seen before), were considered to have gained explicit awareness ^32,33^.

### MRI Acquisition and Data Processing

Structural and functional brain images were acquired using a Philips Elition 3.0 T MRI scanner housed in the Djavad Mowafaghian Centre for Brain Health at the University of British Columbia. First, a T1 structural scan with the following parameters: TR = 8.1ms, TE = 3.61 ms, flip angle = 8°, field of view = 256 × 256, 165 slices, 1 × 1 × 1 mm thickness, was completed. Resting state scans were collected in the same scanning sessions and were acquired using two single shot echo planar imaging sequence with the following parameters: TR = 2000ms, TE = 30ms, flip angle = 90°, voxel dimension = 3 × 3 × 3 mm, 1 mm gap, 36 slices, field of view = 240 × 143 × 240 mm, for a total imaging time of 4.2 minutes for each scan. During the two scans totaling 8.4 minutes, participants were asked to look at a fixed target and to think of nothing in particular.

Resting-state fMRI preprocessing was completed using Statistical Parametric Mapping (SPM 8, University College London, London, UK). A standard pre-processing protocol of realignment, slice-timing correction, outlier detection, segmentation and normalization were used. The realigned functional images were warped into the normalized Montreal Neurological Institute (MNI) EPI template, resampled into 2 × 2 × 2 mm^3^ resolution. Finally, the normalized images were smoothed with a Gaussian kernel at 6mm.

### Functional Connectivity Analysis

Analysis of functional network connectivity was carried out using CONN v.19c ^34^, a functional connectivity toolbox. The white matter and cerebrospinal fluid time series were decomposed into principal components using the CompCor method ^35^, which were regressed out of the total fMRI signal. Five principal components were used for the white matter and the cerebrospinal fluid. Residual head movement parameters (six for rotation and translation parameters, and another six parameters representing their first-order temporal derivatives, outliers identified by ART) were regressed out before analysis. Time points with excessive motion (>.9 mm), or where a global signal changed by above five standard deviations, were defined as outliers and “scrubbed,” and removed from analysis. Resting-state data were band-pass filtered (0.008−.09 Hz).

### Independent Component Analysis

After denoising, group-level independent component analysis (ICA) was implemented across all participants through Calhoun’s group-level fast ICA approach which first performs the ICA, then those components are back projected to individual subject ICA maps which are then entered into the second-level analysis ^36^. The number of independent components was set to 20 ^37,38^ and dimensionality reduction was set to 64 to detect resting-state networks ^36^. Independent components corresponding to resting-state networks were visually identified and confirmed with the spatial correlation of the independent components with the CONN functional atlas. Each group-level spatial map was compared to CONN’s default networks file. Components that were identified as representing intrinsic networks had correlation coefficients between *r* = 0.3 and *r* = 0.5. Thus, the overlap between CONN’s default network maps and the ICA produced relatively high correlation coefficients. A contrast for intervention group ([−1 1]) and time point ([1 −1]) were performed on each ICA to understand network differences between the rest and exercise group between pre and post scanning sessions. Voxel threshold was uncorrected *p* < .001 and cluster threshold was FWE corrected *p* < .05.

### Seed-to-voxel Analysis (regions of interest)

As previous results have demonstrated changes in functional connectivity in brain regions associated with the DMN and limbic system after exercise ^19,39^, we chose 6 brain regions of interest (ROIs) that comprise of the limbic system and were available in the CONN toolbox. The CONN toolbox uses parcellations derived from the Harvard-Oxford Cortical atlas for cortical and subcortical regions ^40^ and the AAL atlas for cerebellar areas ^41^. The ROIs chosen included bilateral anterior division of parahippocampal gyrus, bilateral posterior division of parahippocampal gyrus, bilateral hippocampus, bilateral amygdala. In a follow-up analysis, we also included an additional 14 cerebellar ROIs, which included bilateral lobules 1-10 based on initial seed-to-voxel results from the hippocampus and amygdala. First-level analysis was performed using a weighted general linear model, hemodynamic response function weighting, and bivariate correlation, CONN’s default. For second-level analysis, a voxel threshold was performed at an uncorrected level (*p* < .001) and a family wise error (FWE) correction was applied at the cluster level (*p* < .05).

### Data Analysis

All statistical analyses were carried out using SPSS software (SPSS 19.0; IBM Corporation, Armonk, NY). All data was tested for normality with the Shapiro-Wilk test with alpha .001 ^42^. A Mann-Whitney U (non-parametric test) was run to assess differences in explicit awareness.

Similar to Boraxbekk et al. (2016), each participant’s physical fitness was calculated as a composite z-score taken from five measures (blood pressure: diastolic and systolic at rest, resting heart rate, Godin, and max watts achieved during the stress test) collected at baseline. Each variable was separately z-transformed and then averaged into a composite z-score. Variables with higher values that may indicate poorer physical fitness (diastolic, systolic, and resting heart rate) were multiplied by −1. Thus, the greater positive value of the composite z-score indicated greater physical fitness. The composite score was used as a covariate for all analyses.

Reaction time (RT), the time between movement onset after the start of a trial was used to measure motor acquisition and learning. Additionally, cumulative magnitude (CM), the sum of the square root (X^2^ + Y^2^), where X^2^ and Y^2^ is the absolute value of the derivative of all × and Y coordinates within a trial of the moving hand, respectively was used to understand whether exercise affected movement efficiency. The first trial (always comprised of 7 reaches and random) as well as any reaches equal to or exceeding 1000 ms in reaction time and their neighboring reaches that comprised the same trial were removed from analysis. Values for RT and CM were obtained for each target and movement and were averaged across 7 movements for the random sequence or 6 movements for the repeated sequence.

Averaged trial RT and CM were then calculated separately as ratios relative to the averaged pre-test for the corresponding trial sequence type (e.g., averaged pre-test random trial / first random trial in practice). Thus, higher ratios indicated greater change during acquisition and characterized motor learning at the retention tests. Ratios for each sequence type were averaged across the four blocks of practice for each day for each participant. The ratios from each sequence type were used in three separate mixed-model analysis of variance (ANOVA)(one for RT and CM) with target type (random, repeated) and day (practice day 1-5) being within-subjects factors and group (exercise, rest) as the between-subjects factor. Pairwise post-hoc analyses were Bonferroni adjusted. Separate one-way repeated measures ANOVAs were used to understand group differences in motor learning for the short- and long-term retention tests.

## Results

Of the 44 participants recruited, two participants’ behavioural data were omitted from analysis: one was discarded because of technical issues that occurred during the motor learning task, and data from another participant was discarded because of outliers (3 SD from the mean). One participant from the exercise group and one participant from the rest group were classified as moderately active (14-23) and 2 participants from the rest group were considered sedentary (<14)(Godin, 2011). All other participants were classified as active (≥24 on Godin). All participants stayed within their respective physical fitness levels (sedentary, moderately active, physically active) when assessed at the post-intervention time point. A sum of the raw Godin scores and participant demographics are reported in Table 1. Heart rate and RPE scores averaged across exercise periods (e.g., warm-up, high-intensity period 1, active recovery period 1, etc.) for the exercise group are displayed in Table 2. All participants completed the rest or exercise protocol with no adverse effects. One participant from the exercise group did not undergo the post-intervention resting state fMRI (rs-fMRI) scans due to claustrophobia. Two participants did not come back for the 35-day retention due to scheduling conflicts and 5 participants (3 from exercise, 2 from rest) were excluded from 35-day retention analysis as the number of days between the 24-hour retention and 35-day retention was greater than 2 standard deviations from the mean. A repeated measures ANOVA was used to determine whether there were differences in the number of days between practice sessions. On average there were 3.76 days between each practice session (exercise group range: 2-7, rest group range: 2-6 days) but no main effect of time, no differences between the groups, and no day by group interaction. There was an average of 34.0 days between the 24-hour retention and 35-day retention (exercise group range: 26-39, rest group range: 28-48 days). There was no difference between the number of days between the 24-hour retention session and the 35-day retention session for the two groups.

**Table 2.** Mean heart rate (SD) and rate of perceived exertion (RPE) averaged across 5 day intervention for the exercise group.

| **Period** | **Heart Rate** | **RPE** |
| --- | --- | --- |
| Rest | 73.5 (10.6) |  |
| Warm-up | 84.7 (12.6) | 1.3 (.8) |
| HITT 1 | 114.1 (17.8) | 4.3 (1.7) |
| Recovery 1 | 102.7 (94.7) | 1.7 (.9) |
| HITT 2 | 117.3 (21.1) | 4.6 (1.5) |
| Recovery 2 | 95.8 (15.6) | 1.8 (.9) |
| HITT 3 | 120.5 (24.0) | 4.7 (1.7) |
| Recovery 3 | 98.8 (16.7) | 1.7 (1.0) |

### Resting-State Functional Connectivity (rs-fMRI): Exercise greater than rest, post greater than pre

#### Functional Connectivity (ICA) Networks Group Effects

Of the twenty components, three showed significant increases in the exercise group relative to the rest group. The exercise group displayed greater functional connectivity changes between cerebellar, frontal-parietal, and dorsal attentional networks and bilateral putamen (Table 4, top). Relative to the rest group and the pre time point, the exercise group showed increased connectivity from the cerebellar network to the left putamen (Figure 3a). Similarly, the exercise group displayed an increase in FC between the frontoparietal network and the left frontal pole and left middle frontal gyrus (Figure 3b) and between the dorsal attentional network and right putamen and right insular cortex (Figure 3c).

**Table 3.** Mean reaction time (SD) for rest and exercise groups across sessions.

|  | Rest |  | Exercise |  |
| --- | --- | --- | --- | --- |
|  | Random | Sequence | Random | Sequence |
| Day 1 | 0.411 (.02) | 0.397 (.01) | 0.407 (.02) | 0.396 (.01) |
| Day 2 | 0.400 (.01) | 0.382 (.01) | 0.391 (.01) | 0.374 (.01) |
| Day 3 | 0.389 (.01) | 0.369 (.01) | 0.383 (.01) | 0.364 (.01) |
| Day 4 | 0.381 (.01) | 0.357 (.01) | 0.381 (.01) | 0.358 (.01) |
| Day 5 | 0.379 (.01) | 0.357 (.01) | 0.381 (.01) | 0.359 (.01) |
| 24-hour | 0.370 (.01) | 0.346 (.01) | 0.378 (.01) | 0.355 (.01) |
| 30-day | 0.393 (.01) | 0.374 (.01) | 0.388 (.01) | 0.368 (.01) |

**Table 4.** Significant ICA and seed-to-voxel positive contrasts exercise > rest, post > pre.

|  | *k* | p-FWE | T | Clusters<br>(x,y,z) | Region |
| --- | --- | --- | --- | --- | --- |
| **ICA** |  |  |  |  |  |
| Cerebellum | 130 | 0.015 | 5.57 | -18 -2 -22 | L putamen, L Amygdala |
| Frontoparietal | 174 | 0.002 | 6.27 | -38 38 24 | L FP, L MFG |
| Dorsal attentional | 194 | <0.001 | 7.56 | 22 24 -8 | R putamen, R IC |
| **Seed-to-voxel** |  |  |  |  |  |
| L hippocampus | 95 | 0.04 | 4.76 | 18 -72 -24 | R crus1, R lobule 6 |
| L amygdala | 254 | <0.001 | 5.21 | 24 -76 -34 | R crus2, R crus1 |
| R lobule 9 | 132 | 0.011 | 5.58 | 68 -30 -2 | R MTG, R ITG |
| R lobule 10 | 254 | <0.001 | 5.69 | -56 -52 -06 | L MTG, L ITG |
|  | 113 | 0.019 | 5.49 | 64 -44 -8 | R MTG |
|  | 107 | 0.025 | 5.16 | -54 -24 -6 | L MTG, L STG |

FP = frontal pole, FWE = family wise error, IC = insular cortex, ITG = inferior temporal gyrus, *k* = spatial extent (voxel), MFG = middle frontal gyrus, MTG = middle temporal gyrus, STG = superior temporal gyrus

**Figure 3.**
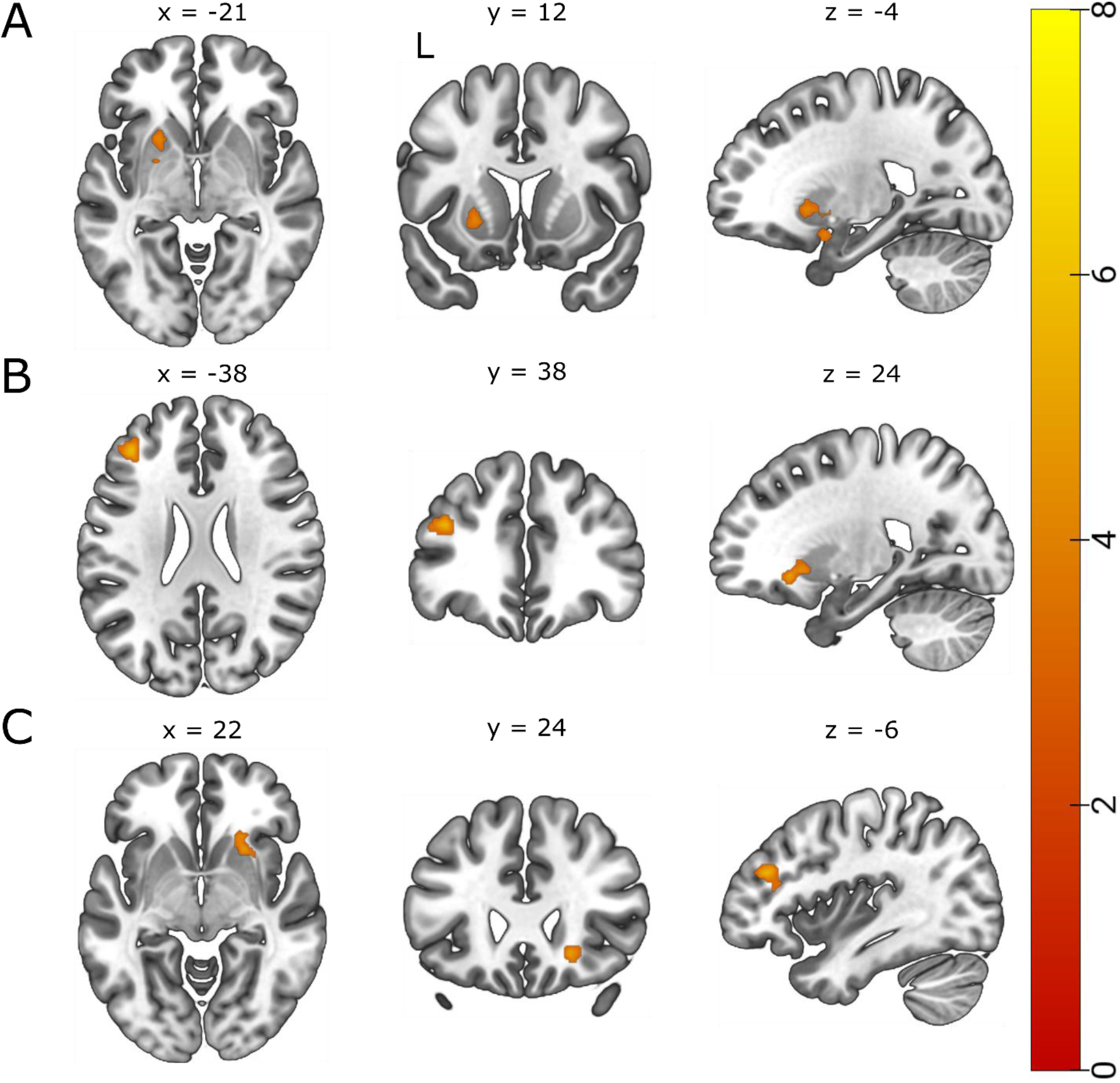
*t* statistical map from ICA exercise > rest, post > pre. Slice views of significant peaks (cluster threshold *p* < 0.05 FWE-corrected, *p* < 0.001 uncorrected at voxel level) that emerged between A) the cerebellar network and left putamen (T(39) > 5.57, k = 130, *p*-FWE = 0.015, B) the frontoparietal network and left frontal pole and middle frontal gyrus (T(39) = 6.27, k = 174, *p*-FWE = 0.002 and C) the dorsal attentional network and right putamen (T(39) = 7.56, k = 194, *p* < 0.001).

#### Functional Connectivity (seed-based) Group Effects

Positive contrasts revealed increased FC between brain regions that link the limbic system and the cerebellum (Table 4, bottom). Specifically, the exercise group also displayed a greater change in FC between the left hippocampus and R crus1 and lobule 6 of the cerebellum (Figure 4 blue-green) and between the left amygdala and the right crus2 and crus1 of the cerebellum (Figure 4 red-yellow).

**Figure 4.**
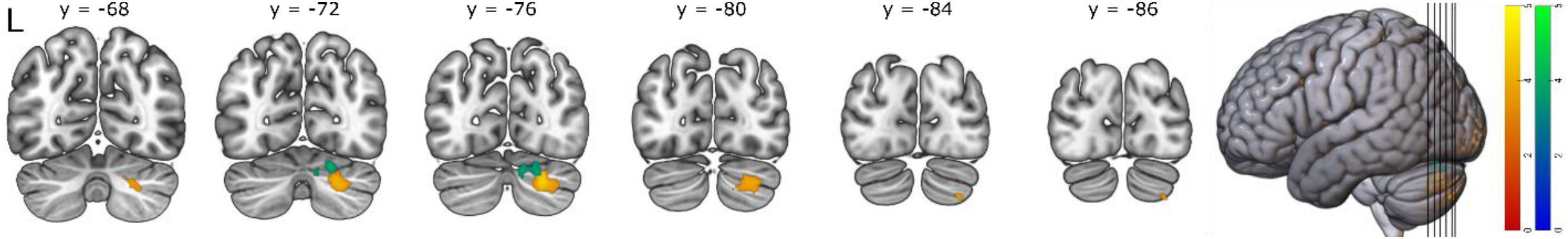
*t* statistical map from seed-to-voxel exercise > rest, post > pre. Slice views of significant peaks (cluster threshold *p* < 0.05 FWE-corrected, *p* < 0.001 uncorrected at voxel level) using the left hippocampus (blue-green) and left amygdala (red-yellow) as seed regions. Increased connectivity observed between the left hippocampus (seed) and the right crus I and lobule 6 of the cerebellum (blue-green)(T = 4.76, *k* = 95, *p*-FWE = 0.040) and between the left amygdala (seed) and the right crus I and II of the cerebellum (red-yellow)(T = 5.21, k = 254, *p*-FWE < 0.001).

To further explore the role of exercise on resting state FC in the cerebellum, we included 14 ROIs. Seed-to-voxel Analysis revealed enhanced FC limited to the right lobule 9 and 10 seed regions of the cerebellum. The exercise group displayed increased FC between the right lobule 9 of the cerebellum and right middle temporal gyrus (Figure 5a), and between the right lobule 10 of the cerebellum and bilateral middle temporal gyri, left inferior temporal gyrus, and left superior temporal gyrus (Figure 5B).

**Figure 5.**
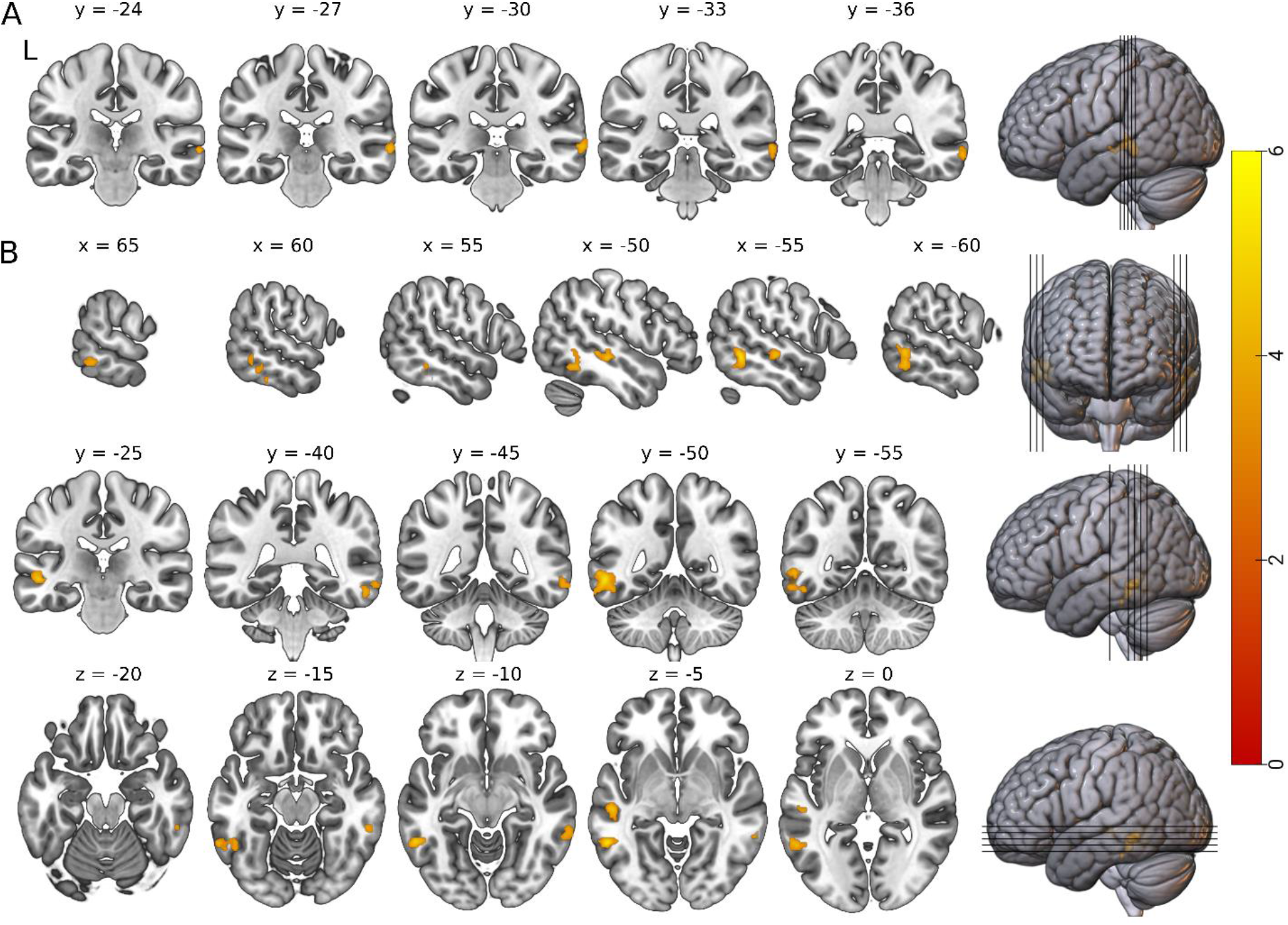
*t* statistical map from seed-to-voxel exercise > rest, post > pre. Slice views of significant peaks (cluster threshold *p* < 0.05 FWE-corrected, *p* < 0.001 uncorrected at voxel level) using A) the right lobule 9 of the cerebellum and B) the right lobule 10 of the cerebellum as the seed regions. Slice views of significant peaks (cluster threshold *p* < .05. cluster size *p* < .05 FWE-corrected) between A) the right lobule 9 of the cerebellum (seed) and the right middle temporal gyrus (T(39) = 5.58, k = 132, *p*-FWE = 0.011), and between B) the lobule 10 of the cerebellum (seed) and three significant clusters. The first cluster was located in the left middle temporal gyrus and the left inferior temporal gyrus (T(39) = 5.69, k = 254, *p*-FWE < 0.001), the second cluster was located in the right middle temporal gyrus (T(39) = 5.49, k = 113, *p*-FWE = 0.019), and a third cluster located in the left middle temporal gyrus and left superior temporal gyrus (T(39) = 5.16, k = 107, *p*-FWE = 0.025).

### Behavioural Data

All dependent variables (RT, CM) were normally distributed for both the exercise and rest groups (p > .001). A total of 276 movements (.3% of total movements) were outliers (1000 ms or greater from the mean). One hundred and twenty-one of those movements (81 random) were from the exercise group and 155 movements (107 random) were from the rest were group.

### Explicit Awareness

Three participants from the exercise group and 9 participants from the rest group acquired explicit knowledge of the repeated sequence when assessed during the 24-hour retention session. Four participants from the exercise group and 10 participants from rest group acquired explicit knowledge of the repeated sequence when assessed during the 35-day retention session. A Mann-Whitney U test revealed no significant difference in explicit awareness between the exercise and the rest group at the short-term (*U* = 156.5, *p* = .08). or long-term retention (*U* = 152, *p* = .22). To understand whether explicit awareness changed between the 24-hours to 35-day retention tests, a change score (35-day minus 24-hour post intervention) was calculated. We found no significant difference in awareness between the rest and exercise groups across time (*U* = 163, *p* = .26).

### Reaction Time

Independent samples t-tests revealed that there was no difference in baseline pre-test RT between the rest and exercise group for the repeated sequence (t(39) = −.737, *p* = .47), or random sequence (t(37) = −.852, *p* = .40).

Using the averaged RT ratio data for each participant as the dependent variable across the 5 days of practice, a 2 (sequence: repeated, random) by 5 (practice day: 1, 2, 3, 4, 5) mixed-model ANOVA revealed a significant main effect of sequence type (*F*(1,38) = 63.819, *p* < .001, η_p_^2^ = .627 and day (*F*(4,152) = 23.410, *p* < .001, η_p_^2^ = .381), but no main effect of group (*F*(1,38) = .684, *p* = .413, η_p_^2^ = .018). Additionally, there was a significant sequence by day interaction (*F*(4,152) = 14.940, *p* < .001, η_p_^2^ = .282). However, there was no sequence by group (*F*(1,38) = .720, *p* = .402, η_p_^2^ = .019), practice day by group (*F*(4,38) = 1.233, *p* = .299, η_p_^2^ = .031), or sequence by practice day by group (*F*(4,38) = .098, *p* = .983, η_p_^2^ = .003) interactions (Figure 6, left). Mean RT values for sequence and random trials can be found in Table 3.

**Figure 6.**
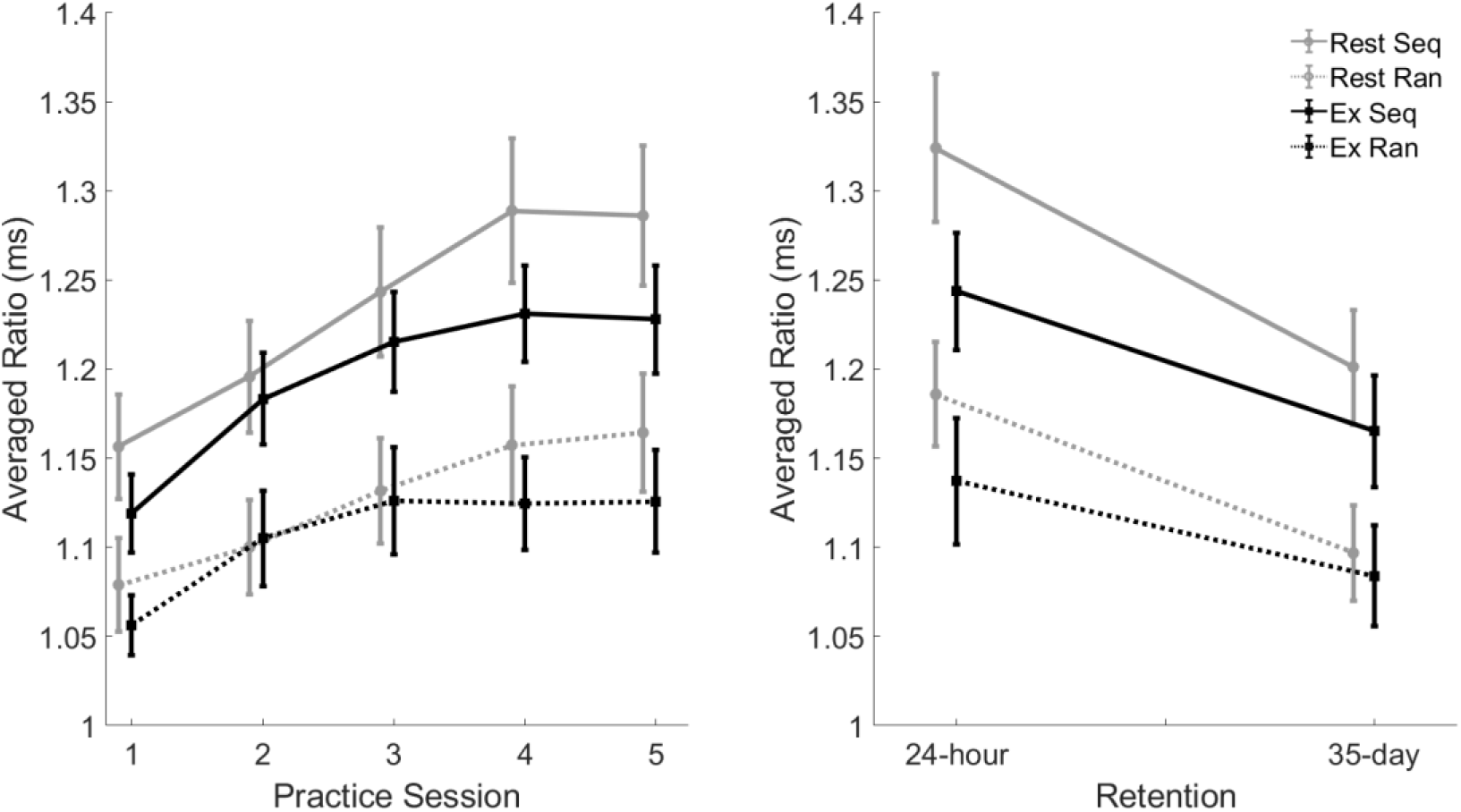
Averaged ratio reaction time (RT) as a function of practice session (left) and 24-hour and 35-day retention session (right) for exercise and rest groups from the serial targeting task. All values were shown as a ratio relative to pre-test, then averaged across four blocks of practice each session. Higher values indicate better motor performance (left) or greater change associated with motor learning (right). Error bars represent standard error. Analysis revealed no main effect of group for practice (*p* = .413), 24-hour (*p* = .184) or 35-day retention (*p* = .793).

Post-hoc comparisons revealed that participants demonstrated greater change in RT across practice for the repeated sequences (M = 1.215, SE ± .021) compared to the random sequences (M = 1.117, SE ± .018; *p* < .001). To understand the main effect of practice day, post-hoc comparisons revealed that participants improved performance across days. Performance change on practice day 2 (M = 1.146, SE ± .019, *p* = .001) was significantly greater than practice day 1 (M = 1.103, SE ± .016). Similarly, performance change on practice day 3 (M = 1.179, SE ± .021) was significantly greater than practice day 2 (*p* = .033). However, there was no significant difference between performance change on practice day 3 and practice day 4 (*p* = .457) and practice day 4 and practice day 5 (*p* = 1.0).

To test motor learning, two separate one-way repeated measure ANOVAs revealed a significant main effect of sequence at the 24-hour retention test (*F*(1,38) = 78.498, *p* < .001, η_p_^2^ = .674), but no main effect of group (*F*(1,38) = 1.834, *p* = .184, η_p_^2^ = .046), and no sequence type by group interaction (*F*(1,145) = 1.475, *p* = .232, η_p_^2^ = .037)(Figure 6, right). Post-hoc comparisons revealed there was more learning related change for in the repeated sequences (M = 1.284, SE ± .027) relative to the random sequences (M = 1.162, SE ± .023, *p* < .001) at the 24-hour retention session.

A similar pattern emerged at the 35-day retention test (Figure 6, right). There was a significant main effect of sequence type (*F*(1,34) = 37.109, *p* < .001, η_p_^2^ = .522), but no main effect of group (*F*(1,34) = .070, *p* = .793, η_p_^2^ = .002) and no sequence type by group interaction (*F*(1,34) <.001, *p* = .997, η_p_^2^ < .001). Post-hoc analysis revealed that participants showed greater learning related change for the repeated sequence (M = 1.128, SE ± .053) relative to the random targets (M = 1.036, SE ± .048, *p* < .001).

Since there were no group differences in RT at the 24-hour and 35-day retention, the groups were collapsed, and individuals were categorized based on explicit awareness of the sequences. There were no group differences between the explicit and implicit learners in the 24-hour retention (*F*(1,38) = .013, *p* = .908, η_p_^2^ < .001) or the 35-day retention (*F*(1,36) = .418, *p* = .522, η_p_^2^ = .011).

### Cumulative Magnitude

A mixed-model ANOVA using the averaged CM ratio for each participant for each day as the dependent variable revealed a significant main effect of sequence (*F*(1,38) = 78.686, *p* < .001, η_p_^2^ = .674) and practice day (*F*(4,152) = .4.652, *p* = .001, η_p_^2^ = .109), but no main effect of group (*F*(1,38) = 1.073, *p* = .307, η_p_^2^ = .027). There was also a significant sequence by practice day interaction (*F*(4,152) = 34.836, *p* < .001, η_p_^2^ = .478). In contrast, there were no significant sequence by group (*F*(1,38) = 1.831, *p* = .184, η_p_^2^ = .046), practice day by group (*F*(4,38) = .778, *p* = .541, η_p_^2^ = .020), or sequence by practice day by group (*F*(4,152) = .315, *p* = .868, η_p_^2^ = .008) interactions.

Follow-up, post-hoc comparisons revealed that across all participants there was greater performance related change for the repeated sequence (M = 1.032, SE ± .007) relative to the random targets (M = .975, SE ± .005, *p* < .001). Regarding the main effect of practice day, there was greater performance related change on practice day 4 (M = 1.008, SE ± .005) relative to practice day 2 (M = .998, SE ± .005, *p* = .007). No other comparisons reached significance.

Within the 24-hour retention session there was a significant main effect of sequence (*F*(1,38) = 23.439, *p* < .001, η_p_^2^ = .382), no main effect group (*F*(1,38) = 1.397, *p* = .245, η_p_^2^ = .035), and no sequence by group interaction (*F*(1,38) = 3.591, *p* = .066, η_p_^2^ = .086). Post-hoc comparisons showed there was greater learning related change for the repeated sequences (M = 1.034, SE ± .006) compared to the random sequences (M = 1.000, SE ± .005, *p* < .001).

At the 35-day retention session, there was significant main effect of sequence (*F*(1,36) = 13.143, *p* = .001, η_p_^2^ = .267), but no main effect of group (*F*(1,36) = .000, *p* = .985, η_p_^2^ = .000), and no sequence or group interaction (*F*(1,36) = .633, *p* = .431, η_p_^2^ = .017). Pairwise comparisons revealed there was significantly greater learning related change for the repeated sequences (M = 1.034, SE ± .006) relative to the random sequences (M = 1.008, SE ± .006, *p* = .007). Collapsing across the rest and exercise group and categorizing individuals based on explicit awareness of the sequences, there was no difference between the explicit and implicit learners at the 24-hour retention (*F*(1,38) = .664, *p* = .420, η_p_^2^ = .017) or the 35-day retention (*F*(1,36) = .060, *p* = .807, η_p_^2^ = .002).

## Discussion

We hypothesized that five days of high-intensity interval exercise paired with skilled motor practice of an implicit sequence task would (1) facilitate motor learning and (2) enhance FC in brain regions that have shown age-related disruptions in older adults. Contrary to our first prediction, we found no advantage in motor learning for the exercise group; both rest and exercise groups demonstrated motor learning. However, in support of our second hypothesis, the exercise group demonstrated increases in resting state FC in the cerebellar, frontoparietal, and dorsal attentional networks. Additionally, seed-based analysis revealed strengthened FC between the limbic system and the cerebellum, and between the cerebellum and middle temporal gyrus in the exercise group.

### Exercise Enhances FC in Critical Brain Regions

Enhanced connectivity was observed among brain regions shown to be affected by exercise. We observed strengthened FC between the left hippocampus and left amygdala and the right cerebellum, between the right cerebellum and middle temporal gyri, and between the cerebellar network and putamen in the exercise group. In animal models, exercise is linked to neurogenesis in the hippocampus ^12,45^, the cerebellum ^46,47^, and the amygdala ^13,14^. In older adults, exercise reduces hippocampal shrinkage ^9,48,49^. In individuals with Parkinson’s disease, exercise temporarily reverses symptoms, likely affecting the striatum ^50,51^. Strengthened FC is thought to be mediated by Hebbian-like processes (i.e., repeated coactivation of brain regions enhances connectivity) ^52–55^. Since previous work showed that these brain regions can be affected following exercise, it follows that we found increased connectivity between these same brain regions following multiple sessions of high-intensity interval exercise.

The seed-to-voxel results revealed right lateralized cerebellar FC increases limited to the exercise group. Specifically, we observed strengthened connectivity between right crus 1, right crus 2, and right lobule 6 and the left hippocampus and left amygdala as well as strengthened FC between right lobule 9, right lobule 10 and bilateral middle temporal gyri. While the cerebellum is broadly involved in motor learning ^56–60^, it is integral for error-based learning ^57,61^. Age-related cortico-cerebellar FC decreases have been previously documented in older adults ^62,63^ with increased cortico-cerebellar connectivity hypothesized to contribute to more efficient sensorimotor transformations ^62,64^. The importance of cerebellar involvement in error-based learning coupled with the widespread increases in cortico-cerebellar FC in the exercise group here suggests that exercise may affect cortico-cerebellar circuitry which may enhance error-based learning ^65,66^. In this context, it is plausible that while exercise enhanced FC related to error-based learning, it had little impact on motor learning of the STT, as this task contained no error feedback. Indeed, when examining the type of motor task, Wanner et al. (2020) reported that exercise had a beneficial effect on both skill acquisition and motor learning, via a delayed retention test, for adaptation tasks, but not sequence-learning tasks. However, in addition to the task, Hübner & Voelcker-Rehage (2017) concluded that the type of exercise, exercise intensity, and the timing of exercise relative to practice may all potentially impact the effect of exercise on motor acquisition and learning in older adults. Therefore, it is possible that a behavioural change would have been noted had we employed a different type of motor task; future work should consider this possibility.

Exercise paired with skilled motor practice of a novel task led to enhanced resting state connectivity in frontoparietal and dorsal attentional networks, encompassing regions involved in spatial transformations and attentional processes during motor learning. Further, we observed enhanced connectivity in the cerebellar network, including cerebellar-hippocampal interactions, which are integral for spatial and temporal processing in humans ^68^ and implicated in motor sequence learning ^69^. We also found increased connectivity between the cerebellum and the amygdala and between the cerebellum and the hippocampus. Given the age-related disruptions in resting state networks ^17,62^, increases in FC suggest that exercise may be an effective intervention for reversing declines in brain networks as we age. Specifically, mounting evidence suggests that the cerebellum acts as a critical structure to maintain normal motor and cognitive functioning during aging ^64,70^. Older adults show widespread decreased connectivity between the right lobule 1-6, 10, and crus 1 of the cerebellum and the putamen ^62^ and decreased connectivity between the cerebellum and the medial temporal lobe ^62,71^. Additionally, the FC increases, which were limited to the right cerebellum in the older adult exercise group, are noteworthy considering that older adults display bilateral cerebellar recruitment during motor tasks (e.g., sequence learning and tapping) relative to young adults ^64^. Thus, the increased couplings between limbic brain regions, the putamen, and the middle temporal gyri and the cerebellum in the exercise group suggest that repeated bouts of exercise paired with learning may contribute to enhanced cerebellar activity that could support motor and cognitive functioning in older individuals ^64^.

### Exercise and Motor Skill Learning

Contrary to our hypothesis, there was no added benefit of exercise on motor learning-related outcomes in older adults; both the exercise and rest groups showed motor learning. This finding suggests that changes in FC alone do not necessarily translate to enhance motor learning beyond practice. We observed changes in FC between the cerebellum and limbic system; however, we did not observe changes between the cerebellum and the cortical motor regions that support sequence specific motor learning. It is well established that regions including primary motor cortex and supplementary motor area (SMA) are critical for the formation of motor plans, including the relay of kinematic transformation to the task specific effector ^72–76^. Increased resting state FC between the cerebellum and both the primary motor cortex and SMA is associated with learning a finger-opposition sequence task _77,78_. Additionally, our previous work showed that high-intensity interval exercise led to a release of cerebellar inhibition ^79^. Other work revealed that cerebellum-motor cortex connectivity is similarly altered by practice of a motor sequence task ^80^. In the current study, we did not observe changes in the cerebellar – motor cortical network. Research in young individuals suggests that modulation of cerebellar – motor circuits may underlie the exercised-induced enhancements in motor learning ^27,65^. Thus, it is possible that aging altered the impact of exercise on cerebellar – motor cortical circuits and as a result it did not impact motor skill learning in the same manner as has been shown in younger individuals.

A recent meta-analysis examining the impact of exercise on motor learning conducted by Wanner et al. (2020) suggests that stronger effects of exercise are noted when this intervention follows skilled motor practice, although recent work showed no effect of exercise on learning even when exercise follows motor practice^81^. Though other work shows that exercise preceding skilled motor practice enhanced motor learning, this past research only considered young healthy individuals and single sessions of practice ^6,7,82^. Future work should consider the timing of exercise in older adults, relative to skilled motor practice, to test how the order of these two interventions impacts motor learning.

### Limitations

The number of days in between practice sessions and between the 24-hour and 35-day retention sessions were not identical across participants. While a fixed schedule would have been ideal, such an approach would have been difficult to implement with the study design. As this study spanned across eleven separate sessions taking approximately 90 minutes per session, the time commitment for each participant was considerable. In conjunction with allowing enough time in between each exercise session to facilitate muscle recovery (allowing participants to maintain the 75% of maximum output during the high-intensity intervals), we opted to accommodate participants based on their availability. Importantly, it is unlikely that any changes in FC observed are due to discrepancies in participant scheduling, as we found no difference in the number of days between practice sessions between groups. In addition, only one volume of exercise was tested, namely 3 sets of 3 minutes of high intensity exercise training (9 minutes of high intensity exercise), so the impact of other volumes, intensity, or frequency of exercise on these outcomes needs further investigation. Despite these issues we show that exercise alters brain FC in healthy older adults. Moreover, it is possible the current results may be attributed to the moderate-to-high physical fitness levels of the participants and therefore do not generalize to all older adults. However, we did use a composite z-score comprised of baseline physical fitness measures as a covariate in the analyses. One limitation that may affect the interpretation of the functional connectivity results is scan length. While we employed two, 4 minute scans comparable to other resting state and exercise studies with similar results^19,39^, having longer scan lengths reduces the number of spurious connections and increases intraclass coefficients^83^. Future studies investigating the impact of repeated bouts of high-intensity exercise on functional connectivity should employ longer scan lengths., Lastly, high-intensity exercise may come with increased risks, especially for sedentary individuals with unknown underlying health conditions. It should be noted, however, that all self-reported healthy older adults recruited in the current study were able to complete the high-intensity exercise intervention without any adverse effects despite a wide range of physical fitness. Therefore, this exercise protocol is feasible and can likely be generalized to a healthy older adult populations.

### Conclusions

We observed that pairing separate bouts of high-intensity interval exercise with skilled motor practice strengthens FC without enhancing motor learning in a healthy older adult population. Enhanced FC in the exercise group was found in networks and brain regions that are susceptible to declines in ageing, especially in the cerebellum, indicating that exercise paired with motor practice could lead to protective effects in networks that support motor and cognitive function. Overall, these results may have implications for the application of exercise in reducing age-related disruptions to FC and aiding in healthy ageing in older adults.

The authors declare no competing financial interests.

## Author contributions

B.G., J.L.N., K.L.C., L.A.B. designed research; B.G., B.C., C.B.J., J.L.N. contributed to data collection; B.G. analyzed data; B.G. wrote first draft of the paper; B.G., J.L.N., S.N.K., L.A.B. wrote the paper; all authors edited the paper.

## Acknowledgements

This study was funded by the Canadian Institutes of Health Research Project Grant (PJT-148535: PI LAB).

